# Human oral microbiome characterization and its association with environmental microbiome revealed by the Earth Microbiome Project

**DOI:** 10.1101/732123

**Authors:** Jinlan Wang, Dandan Li, Jianing Wang, Zheng Zhang

**Affiliations:** Physical Examination Office of Shandong Province, Health Commission of Shandong Province, Jinan 250014, China; National Institute of Biological Sciences, Beijing 102206, China; State Key Laboratory of Microbial Technology, Institute of Microbial Technology, Shandong University, Qingdao 266237, China

**Keywords:** Oral cavity, Microbiome, Earth Microbiome Project, Microbial diversity, Environmental microbiome

## Abstract

The oral cavity is an important window for the microbial communication between environment and the human body. The oral microbiome plays an important role in human health. Here, we analyzed 447 datasets from human oral samples published by the Earth Microbiome Project (EMP). The microbes in these human oral samples were taxonomically assigned to at least 266 genera of 18 bacterial and archaeal phyla. Among them, 11 genera with the relative abundance more than 1% were identified as 5 different bacterial phyla. Compared with 815 samples from human gut, nose/pharynx and skin, the oral microbiome showed significantly lower diversity and possessed fewer unknown species than those of other body parts, and had distinct differences in species composition from other body parts. In addition, the oral microbiome showed significant differences in the populations of different countries, which may be determined by the living environment and lifestyle/dietary habits. Finally, the correlation analysis showed highly similarity between the oral microbiome and the microbiomes of Aerosol (non-saline) and Surface (non-saline), two types of environmental microbial habitats related closely to human. Together, these findings expand our understanding to the human oral microbiome.

## 1. Introduction

The oral cavity is an important place for the delivery and exchange of the substances inside and outside the human body, and also the gateway for the pathogens and toxic substances to invade the body. The microbes found in human oral cavity are collectively referred to as the oral microbiome [1–3]. The complex and variable interaction of oral microbes helps the body to fight against the external undesirable stimuli. However, the imbalances of microbial community in oral cavity can lead to the oral diseases, such as the dental caries, periodontitis, oral mucosal diseases and systemic diseases such as gastrointestinal and neurological diseases [2,4–13]. Therefore, the oral microbiome plays an important role in maintaining the balance between human microbial communities and human health [14,15].

The composition of the oral microbiome is complex, and the expanded Human Oral Microbiome Database (eHOMD) has included 770 microbial species of 230 genera in 16 bacterial and archaeal phyla [16]. Of all the species in this database, 57% are officially named, 13% unnamed but cultivated and 30% are known only as uncultivated phylotypes. There is no difference among the oral, gut, and skin microbiome of the newborn babies, but the composition of their oral microbiomes will change significantly as the age increases and the dentition changes [17]. The differences of microbiomes for the same individual at different time points are significantly lower in the oral cavity than in the gut, skin and other body parts [18]. The effects of the early living environment on shaping oral microbes are much more than genetic factors [19]. In addition, lifestyle habits, social factors, and oral pH value also affect the composition of the oral microbiome [20].

The Earth Microbiome Project (EMP) aims to collect as many of the Earth’s microbial communities as possible in order to promote our understanding to the relationship between the microbes and the environment including plants, animals and humans [21,22]. The first data published by EMP contained 27,751 samples from 97 independent studies representing different environmental types, geographic locations, and chemical reactions [23]. All samples were subjected to DNA extraction and sequencing, and the bacterial and archaeal parts of the entire database were analyzed. Here, we used the 447 datasets from human oral samples published by EMP to study the characteristics of human oral microbiome and its association with environmental microbiome.

## 2. Materials & Methods

### 2.1 Human oral sample data acquisition based on EMP data

EMP developed a unified standard workflow that leveraged existing sample and data reporting standards to allow biomass and metadata collection across diverse environments on Earth [23]. The samples submitted by the global community of microbial ecologists were used to proceed the microbiome analysis. The DNA extraction and 16S rRNA amplicon sequencing were done using EMP standard protocols [24]. The sequence data were error-filtered and trimmed to the length of the shortest sequencing run (90 bp) using the Deblur software [25].

The EMP data contains a total of 97 studies and 27,742 samples which are available at ftp://ftp.microbio.me/emp/release1. We acquired 447 human oral samples from EMP study to analyze their microbial diversity (Supplementary data). These oral samples are from 5 independent studies and include the populations from Italy, Puerto Rico, the United States, and Venezuela [24,26,27]. We also selected 216 gut, 253 nasal/pharyngeal and 346 skin samples from EMP study to proceed the compared analysis [24,26,28,29].

### 2.2 EMP Ontology classification

EMP classified the samples in different environments into the corresponding environmental labels [23]. The EMP Ontology (EMPO) classified the microbial environments (level 3) as free-living or host-associated (level 1) and saline or non-saline (if free-living) or animal or plant (if host-associated) (level 2). A subset containing 10,000 samples was then generated which give equal (as possible) representation across environments (EMPO level 3) and across studies within those environments. In this subset, each samples must have ≥ 5,000 observations in the Deblur 90 bp observation table.

### 2.3 Comparison against reference databases and core diversity analyses

The representation sequences of operational taxonomic units (OTUs) were analyzed by the Ribosomal Database Project (RDP) Classifier algorithm using a confidence threshold of 50% against the Silva 16S rRNA gene database [30,31].

The alpha diversity was computed with the input Deblur 90 bp BIOM table rarefied to 5,000 observations for each sample. The alpha diversity included observed OTUs (number of unique tag sequences), Shannon index (Shannon diversity index), chao1 index (Chao 1 index), and faith’s PD value (Faith’s phylogenetic diversity) [32–34].

The clustering of samples was conducted due to storage conditions by principal coordinate analysis (PCoA), based on Bray-Curtis similarity distance. The UPGMA (Unweighted Pair Group Method with Arithmetic Mean) clustering was based on Bray-Curtis similarity distance. Bootstrapping with 1,000 resamplings was performed to determine the robustness of the clustering. All these analyses were performed with the statistical software PAST [35].

Significant difference was evaluated using analysis of variance (ANOVA) by the software package IBM SPSS Statistics.

## 3. Results and discussion

### 3.1 Human oral microbiome composition

We analyzed the sequenced data of the human 447 oral samples from 5 independent studies (Supplementary data). The length of each sequenced 16S rRNA from all the samples was truncated to 90 bp, and then 5,000 observed sequences were randomly extracted from each sample for further calculation. All the samples were removed wrong sequences using Deblur algorithm and calculated the OTUs at the level of single nucleotide accuracy.

The results showed that the average of observed bacterial and archaeal OTUs was 73.81±29.26 in human oral samples, with the maximum of 279 OTUs and the minimum of 25 OTUs in a single sample. The Chao1 index is relatively sensitive to low-abundance species. The average of Chao1 index for human oral samples was 89.20±36.98, ranging from 28.75 to 302.02. The Shannon index can simultaneously reflect the species diversity and the community uniformity. The average of Shannon index for human oral samples was 3.64±0.77, varying from 0.97 to 6.16. The Faith’s PD value is a good measure of phylogenetic diversity, and the average of Faith’s PD value for human oral samples was 11.99 ± 3.03, varying between 6.03 and 31.17. These results indicated that the diversity of human oral microbiome was significantly different among individuals.

The taxonomical results of the 16S rRNA gene sequences showed that the microbes in the human oral samples belonged to at least 264 genera of 17 bacterial phyla and 2 genera of 1 archaeal phylum. The predominant phyla of the human oral microbiome were *Firmicutes, Proteobacteria*, *Bacteroidetes*, *Fusobacteria* and *Actinobacteria*, with the average relative abundance of 37.81%, 30.11%, 17.76%, 9.07% and 4.75%, respectively. In addition, 0.02% of the sequences cannot be classified at the phylum level (Fig. 1A). The total relative abundance of the predominant 11 genera (>1%) were 87.59%, and *Streptococcus* (23.52%), *Neisseria* (15.30%) as well as *Haemophilus* (13.21%) were the top three of the average relative abundance among these 11 genera. Meanwhile, 4.80% of the sequences were not classified at the genus level (Fig. 1B). These 11 high abundance genera in human oral samples were distributed among multiple bacterial phyla, 3 of them were the *Firmicutes*, 2 of them were the *Proteobacteria*, 2 of them were the *Bacteroidetes*, 2 of them were the *Fusobacteria*, and 2 of them were the *Actinobacteria*. Therefore, the human oral microbiome also has a high complexity in species composition.

**Fig. 1.**
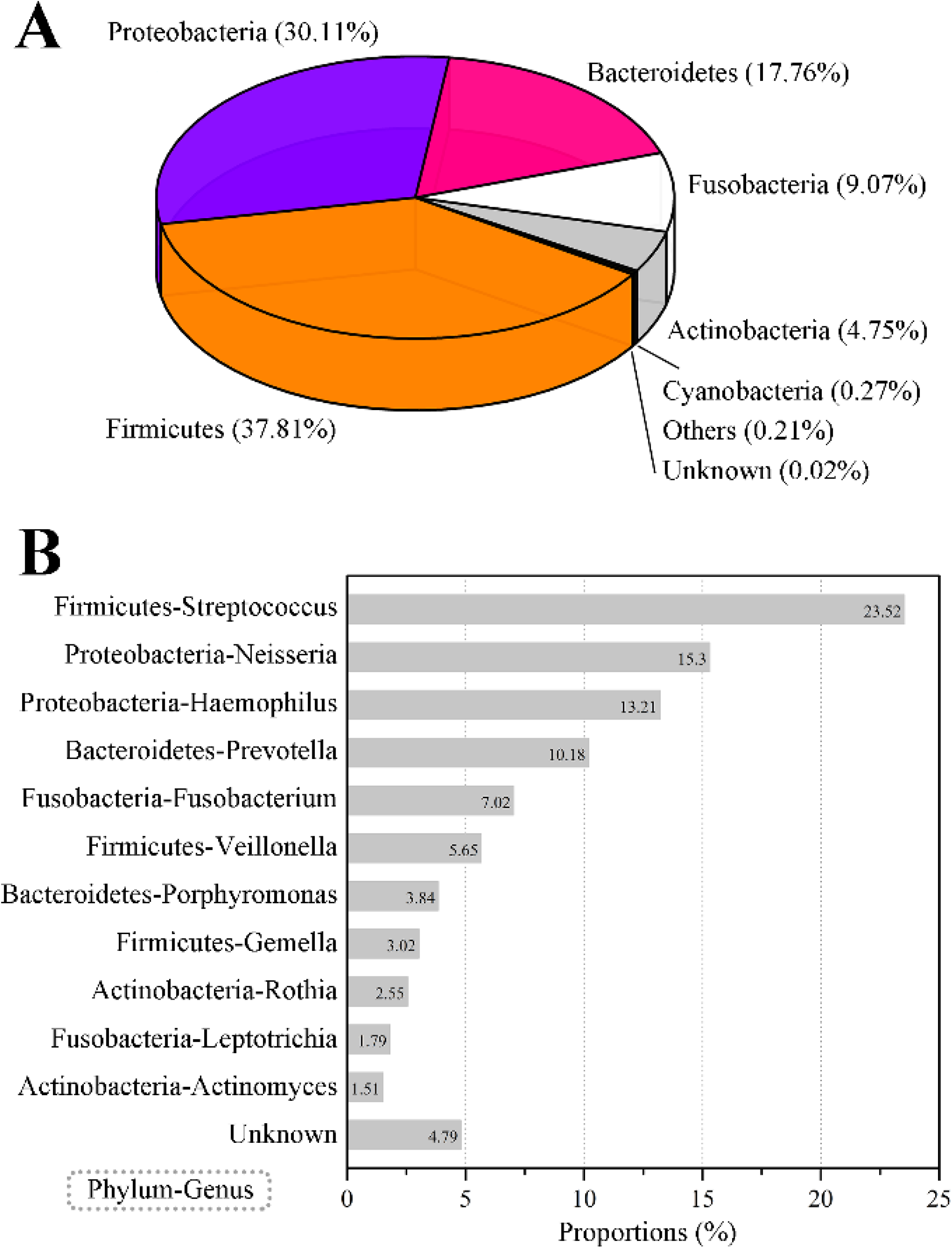
Species composition of human oral microbiome. (A) Species composition of human oral microbiome in phylum level. (B) Eleven human oral microbial genera with larger than 1% abundance. The results were computed based on 447 oral samples of EMP.

### 3.2 Composition comparison of human oral microbes with gut, nasal/pharyngeal and skin microbes

In addition to the oral cavity, the gut, nose/pharynx and skin are also important habitats for human microbial colonization. Using the data published by EMP, we compared the differences between the human oral microbiome and the gut, nasal/pharyngeal, and skin microbiome. Among them, gut microbial data were from 216 samples, nasal/pharyngeal data were from 253 samples, and skin data were from 346 samples.

The results showed that the human oral microbiome diversity was significantly (ANOVA, p < 0.01) lower than that of the gut, nasal/pharyngeal, and skin microbiome (Supplementary Fig. 1). The average of observed bacterial and archaeal OTUs was 116.72±39.64 in human gut samples, 288.76±284.66 in human nasal/pharyngeal samples, and 296.98±176.87 in human skin samples, each of which was significantly higher than the average value observed in human oral samples. The average values of Chao1 index for human gut, nasal/pharyngeal and skin samples were 140.29±49.75, 449.81±469.85, and 422.42±271.39, respectively, which were significantly higher than 89.20±36.98 for oral samples. While, the average Shannon indexes for human gut, nasal/pharyngeal and skin samples were 4.45±0.81, 4.27±2.00, 4.85±1.60, respectively, which were significantly higher than 3.64±0.77 for the oral sample. Besides, the average Faith’s PD values for human gut, nasal/pharyngeal and skin samples were 15.30±4.35, 30.26±21.22, 31.30±14.53, which were also significantly higher than 11.99±3.03 for oral samples.

The results of taxonomy showed that the microbes in human gut samples were at least 288 genera of 16 bacterial phyla and 5 genera of 2 archaeal phyla, slightly more than those of oral samples. However, the microbes in the nasal/pharyngeal samples were at least 832 genera of 32 bacterial phyla and 11 genera of 3 archaeal phyla. The microbes in the skin samples were at least 948 genera of 32 bacterial phyla and 18 genera of 3 archaeal phyla. Their microbial diversities are much larger than that of oral samples.

*Firmicutes* was not only the highest abundance microbe at the phylum level in the oral microbiome, but also had more than 30% abundance in other body locations, and especially its abundance in the gut microbiome was as high as 49.41% (Fig. 2A). *Proteobacteria* had an abundance of more than 25% in the oral, nasal/pharyngeal and skin microbiome, but only 3.99% in the gut microbiome. *Bacteroidetes* accounted for 17.76% in the oral microbiome and 37.35% in the gut microbiome, but only 4.65% and 6.25% in the nasal/pharyngeal and skin microbiome. The abundance of *Fusobacteria* in the oral (9.07%) microbiome was significantly higher than that in the gut (0.84%), nasal/pharyngeal (0.52%) and skin (2.58%) microbiome. The abundance of *Actinobacteria* in the oral (4.75%) microbiome was close to that of the gut (5.08%) microbiome, but significantly lower than that of the nasal/pharyngeal (16.57%) and skin (18.44%) microbiome. The abundance of *Cyanobacteria* in the oral (0.27%) microbiome was higher than that in the gut (0.02%) microbiome, but lower than that in the nasal/pharyngeal (2.52%) and skin (4.17%) microbiomes. Finally, only 0.02% of the 16S rRNA sequences was not classified at the phyla level in the oral microbiome, but the 16S rRNA sequences that could not be classified at the phyla level are much more in the gut (1.89%), nasal/pharyngeal (0.41%), and skin (0.26%) microbiomes.

**Fig. 2.**
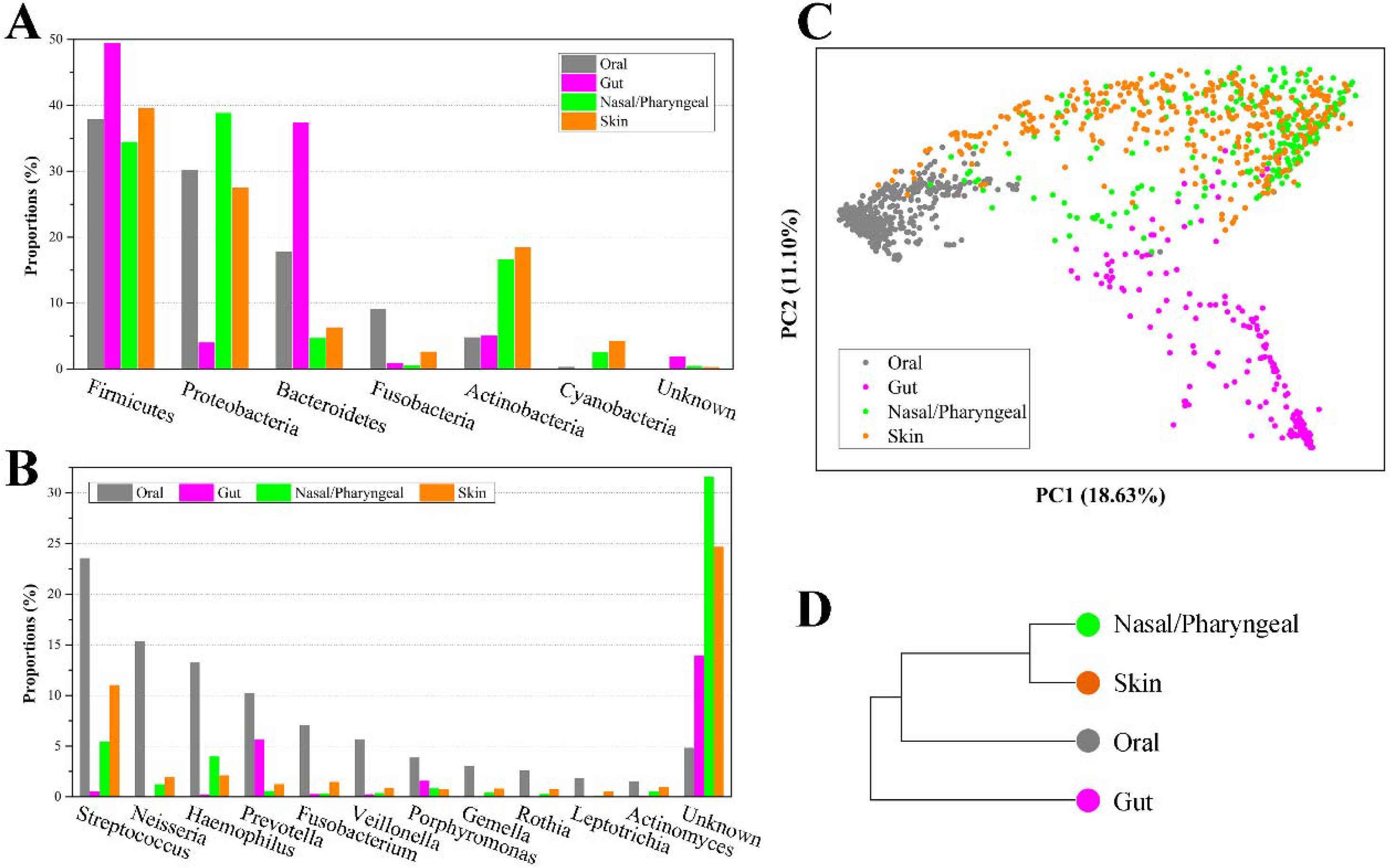
Composition comparisons of human oral cavity, gut, nasal/pharyngeal and skin microbiomes. Distributions of six microbial phyla with the highest abundance (A) and 11 genera with the highest abundance (B) of the oral microbiome at other body parts. PCoA (C) and Clustering (D) analyses of microbial composition from four different parts of human oral cavity, gut, nose/pharynx and skin.

For the 11 genera with more than 1% abundance in the oral microbiome, their abundances in the gut, nasal/pharyngeal and skin microbiomes were significantly lower than that in the oral cavity (Fig. 2B). For example, the abundance of *Neisseria* in the oral cavity was 15.30%, while the abundances in the gut, nasal/pharyngeal and skin microbiomes were only 0.003%, 1.20% and 1.94%, respectively. In addition, the 16S rRNA sequences that couldn’t be classified at the genus level were also largely less in the oral (4.80%) microbiome than in the gut (13.92%), nasal/pharyngeal (31.56%) and skin (24.64%) microbiomes.

Further, we performed a PCoA analysis based on the Bray-Curtis distance for 1,262 samples from human oral cavity, gut, nose/pharynx, and skin, and displayed them in a scatter plot (Fig. 2C). The results showed that the microbiome of oral samples could be well distinguished from the microbiomes of other body parts samples, indicating that the oral microbiome was significantly different from other parts in species composition. Similarly, the microbiome for gut samples could also be well distinguished from the microbiomes of other body part samples. However, there was a lot of overlap for the microbiomes between the nasal/pharyngeal and skin samples. The lowest dispersion of the oral microbiome among the four microbiomes suggested the lowest diversity, which was consistent with the alpha diversity index. The clustering analysis indicated that the nasal/pharyngeal and skin microbiomes were most similar, while the oral microbiome was more similar to the nasal/pharyngeal and skin microbiomes relative to the gut microbiome (Fig. 2D).

In summary, the above results showed that the human oral microbiome was obviously different from the gut, nasal/pharyngeal, and skin microbiomes. The oral microbiome had lower species diversity, fewer unknown species and different species composition from other body location microbes.

### 3.3 Comparison of oral microbiome compositions among different countries

The human oral microbial samples were obtained from 4 countries, including 56 in Italy, 79 in Puerto Rico, 270 in the United States, and 42 in Venezuela. We found that the diversity of the oral microbiome was significantly different (ANOVA, p < 0.01) among the populations of these four countries (Supplementary Fig. 2). For the observed bacterial and archaeal OTUs in the oral cavity, the samples from Italy (106.93±25.04) and Venezuela (93.57±31.12) were significantly higher than those of Puerto Rico (67.84±30.12) and the United States (65.61±22.65). Similarly, the results of Chao1 index, Shannon index, and Faith’s PD value also had the same feature.

In terms of species composition for the Italian samples, the abundances of the *Rothia*, *Leptotrichia* and *Actinomyces* were obviously higher than the average value of each genus of them within 4 countries (the population mean), while the abundances of the *Streptococcus* and *Haemophilus* were lower than the population mean (Fig. 3A). In the Puerto Rican samples, the abundance of *Streptococcus* was increased, while the abundances of *Neisseria*, *Prevotella*, *Veillonella*, and *Porphyromonas* were decreased. In the United States samples, the abundances of *Neisseria*, *Fusobacterium*, and *Oribacterium* were increased, and the abundances of *Streptococcus* and *Gemella* were decreased. In the Venezuelan samples, the abundances of *Leptotrichia* and *Granulicatella* were increased, and the abundance of *Haemophilus*, *Prevotella* and *Veillonella* were decreased. The PCoA analysis based on Bray-Curtis distances showed that the samples from the United States were well distinguished from other countries, whereas the samples from Venezuela partially overlapped with the samples from Italy and Puerto Rico (Fig. 3B). The clustering analysis showed that the samples from Italy were the most similar in microbial composition to the samples from Venezuela, followed by the samples from Puerto Rico, while the samples from the United States were the most unique (Fig. 3C).

**Fig. 3.**
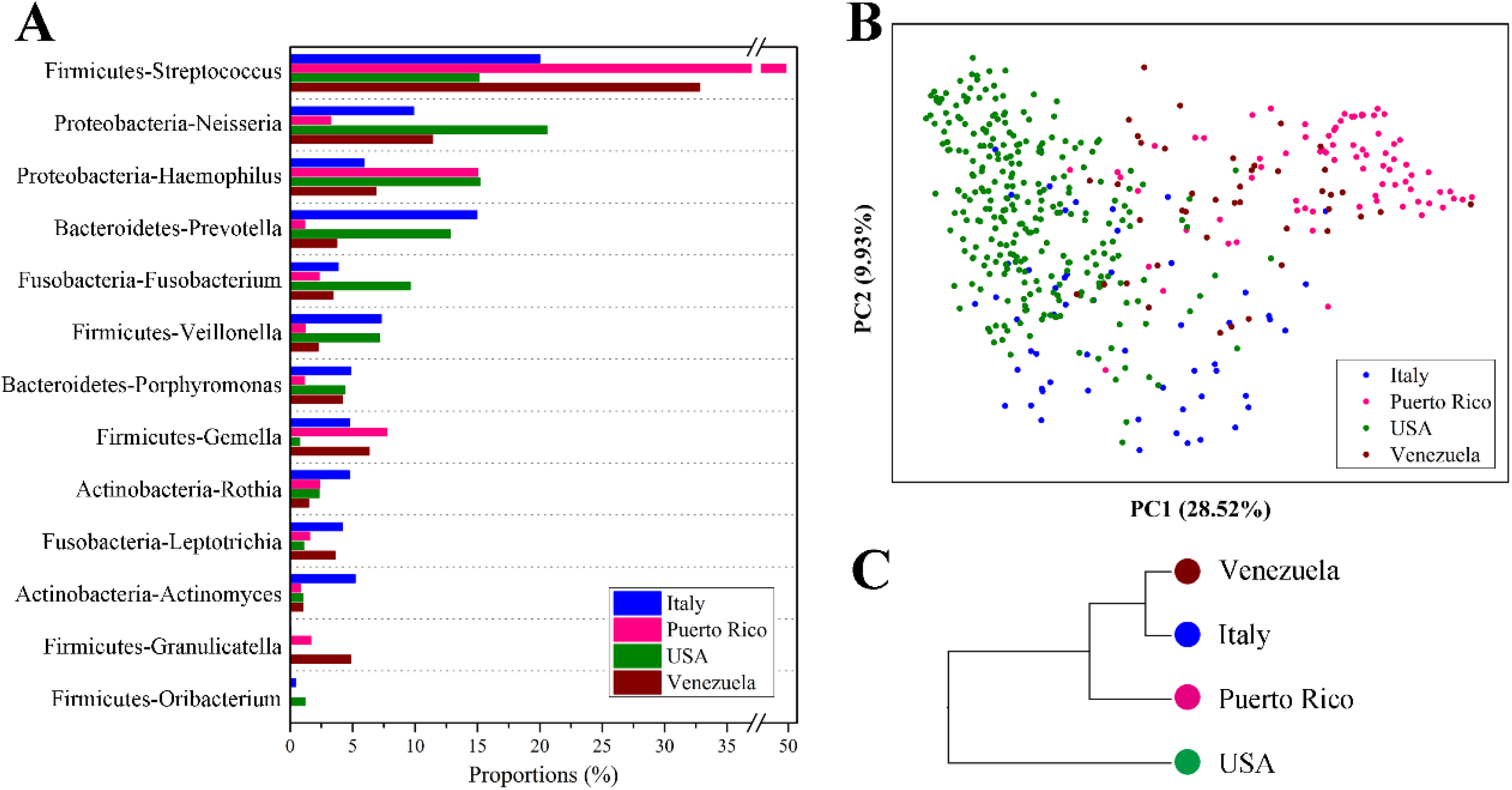
Composition comparisons of the oral microbiomes in four different countries. (A) Comparison of oral microbial composition in populations of four countries. (B) PCoA analysis for the oral microbes from populations in different countries. (C) Clustering analysis of the oral microbes for populations in different countries. Italy, Puerto Rico, the United States, and Venezuela were analyzed and represented by blue, light red, green, and dark red, respectively. A total of 447 samples were used, and the analysis was conducted based on the Bray-Curtis distance.

Finally, we compared the microbial diversity differences in oral, gut, nose/pharynx, and skin across the countries using the samples from the United States and Venezuela. Each country has more than 20 samples for each body location. For the observed bacterial and archaeal OTUs, the oral and gut samples in the United States population were significantly lower (ANOVA, p<0.01) than those in Venezuela, the nasal/pharyngeal samples in the United States population were significantly (ANOVA, p<0.01) higher than that in Venezuelan, whereas the skin samples showed no significantly (ANOVA, p>0.05) differences between the two countries (Supplementary Fig. 3). Similarly, the results of Chao1 index, Shannon index, and Faith’s PD value also showed the same feature. These results indicated that the population, with highly oral microbial diversity, were not always possessing higher microbial diversity of other body parts than the rest of the population.

### 3.4 Association of human oral microbes with environmental microbes

A number of microbes were exchanged with the external environment through human oral cavity. Therefore, we tried to further analyze the association between oral microbes and environmental microbes. EMP has classified the samples in different environments into the corresponding environmental labels. These environmental labels were first divided into two categories: Free-living and Host-associated, and further subdivided into 17 subcategories denominated as EMPO level 3. We performed a cluster analysis to display the association of microbe compositions between human oral, gut, nasal/pharyngeal and skin samples and EMPO environmental labels (Fig. 4). The results showed that the closest EMPO environmental label to human oral samples was Animal secretion, the closest one to human nasal/pharyngeal and skin samples was the Animal surface, and the closest one to human gut samples was the Animal distal gut. Further, the EMPO environmental labels that were close to human oral samples mostly belong to the host-associated type, but also included the two free-living environments of non-saline Aerosol and Surface. Aerosol and surface are the two type environments in the closest contact with human. To be specific, Aerosol is the aerosolized dust or liquid. Surface is the biofilm from wet (<5 psu) or dry surface, wood, dust, and microbial mat.

**Fig. 4.**
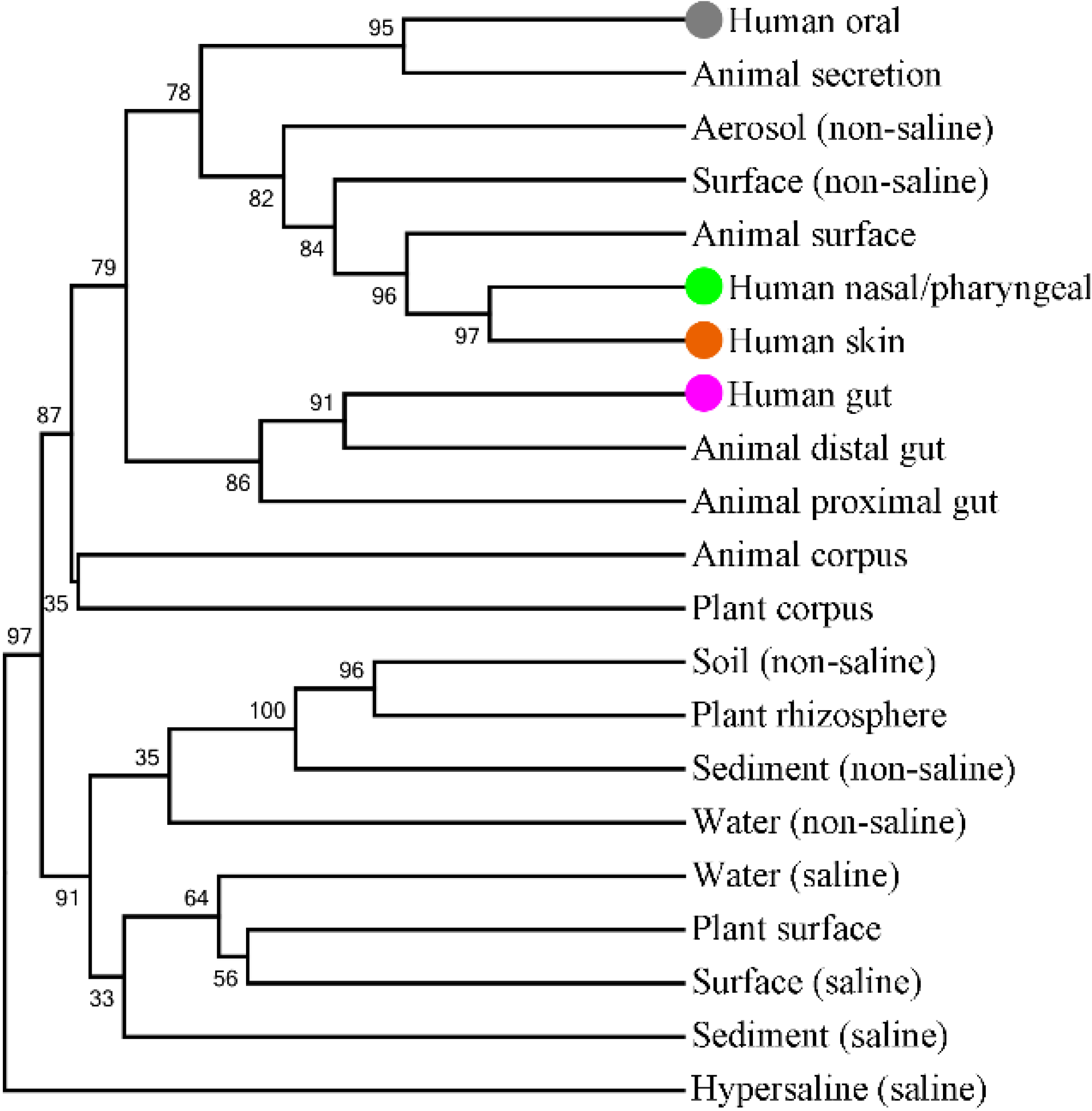
Clustering analysis of microbial composition for human oral, gut, nose/pharynx and skin samples and EMPO environmental labels. The tree was built using the UPGMA method, and Bray-Curtis similarity distance was used. Bootstrapping with 1,000 resamplings was performed to determine the robustness of the clustering. The oral cavity, nose/pharynx, skin and gut were highlighted by grey, light green, orange-red and purple solid spheres, respectively.

For the 11 genera with more than 1% abundance in the oral microbiome, their abundance in all the EMPO environmental labels were obviously lower than those in the oral cavity. For example, the abundance of *Streptococcus* was 23.52% in the oral cavity, 5.07% in Aerosol (non-saline), 4.22% in Animal surface, 3.02% in Surface (non-saline), and 1.35% in Animal proximal gut, and 0.58% in Animal distal gut, and so on. Interestingly, all the 11 genera had the highest abundance in the Aerosol (non-saline) of the nine free-living environments, as well as the second highest abundance in the Surface (non-saline). Further, we found that the abundance of these 11 genera in various environments had an obviously positive correlation. Therefore, the composition of the oral microbes represented by these 11 genera was specific and had a certain similarity with the microbial composition in the free-living Aerosol and Surface environments.

## Conflicts of interest

The authors declare that they have no conflict of interest.

## Acknowledgements

This work was supported by China Postdoctoral Science Foundation (2018M642649 and 2019T120586).

