## Supplementary figure 1-3 for "Human oral microbiome characterization and its association with environmental microbiome revealed by the Earth Microbiome Project"

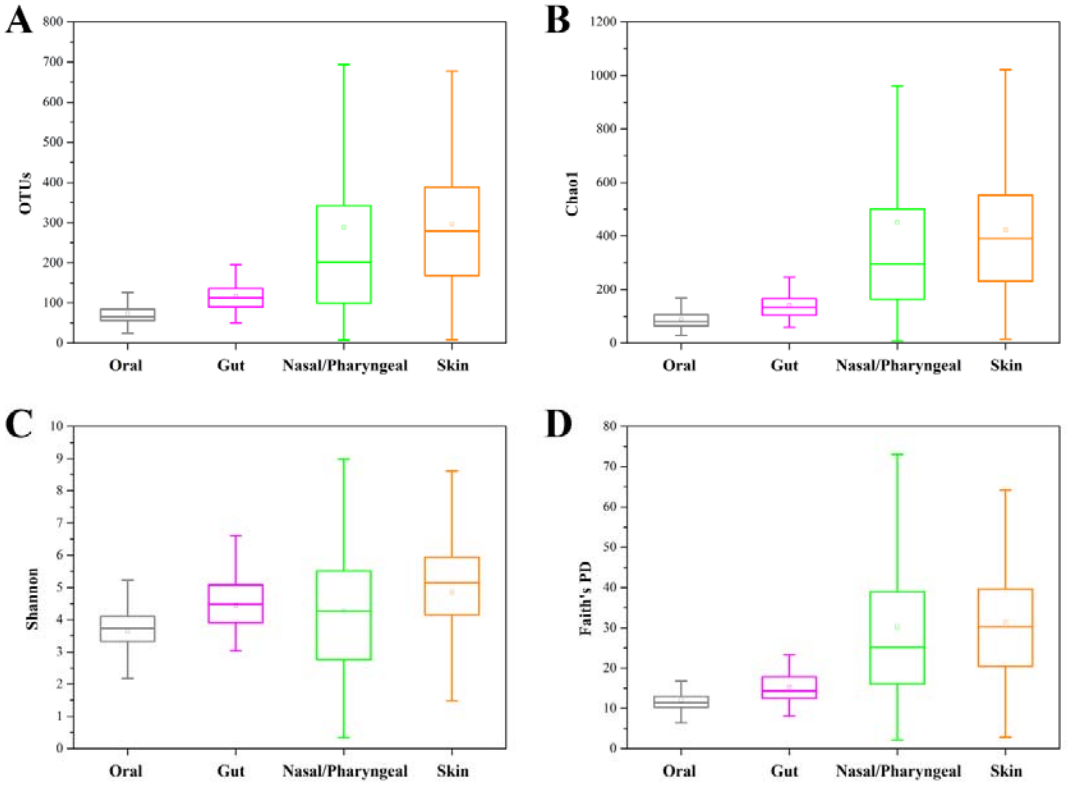


**Supplementary Fig. 1. Alpha diversity comparisons of human oral cavity, gut, nasal/pharyngeal and skin microbiomes**. (A) Observed OTUs. (B) Shannon index. (C) Chao1 index. (D) Faith's PD value.

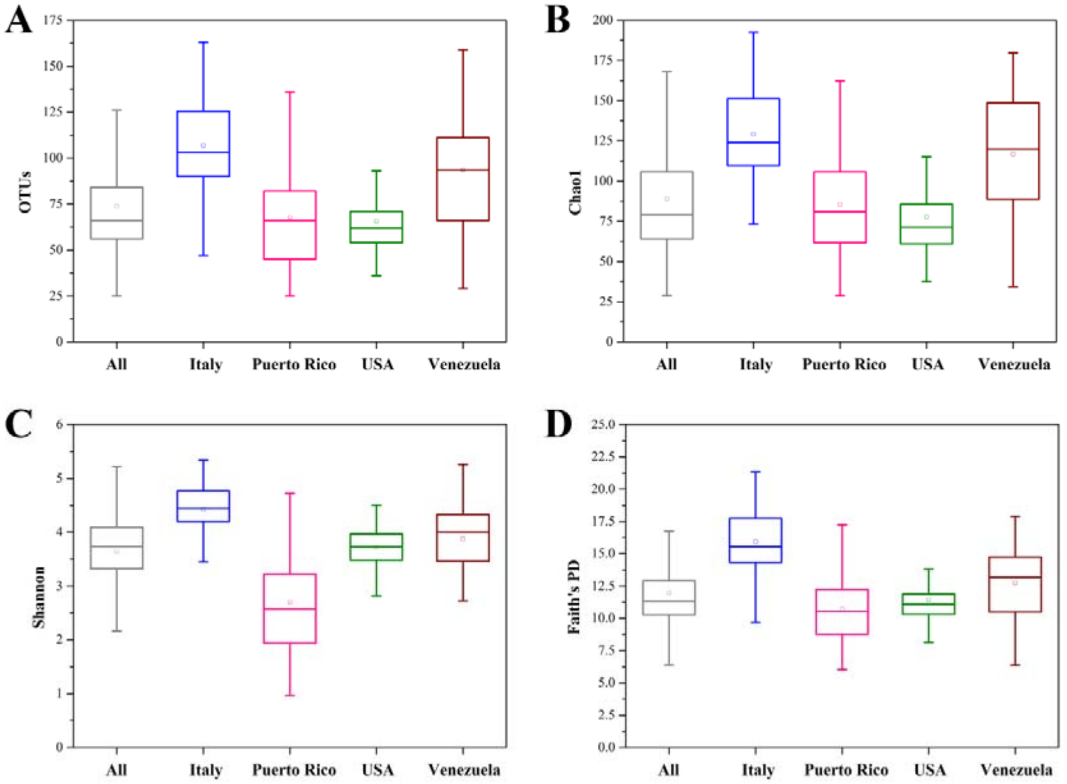


**Supplementary Fig. 2. Alpha diversity comparisons of oral microbiomes in populations from four different countries of the United States, Italy, Puerto Rico, and Venezuela.** (A) Observed OTUs. (B) Shannon index. (C) Chao1 index. (D) Faith's PD value.

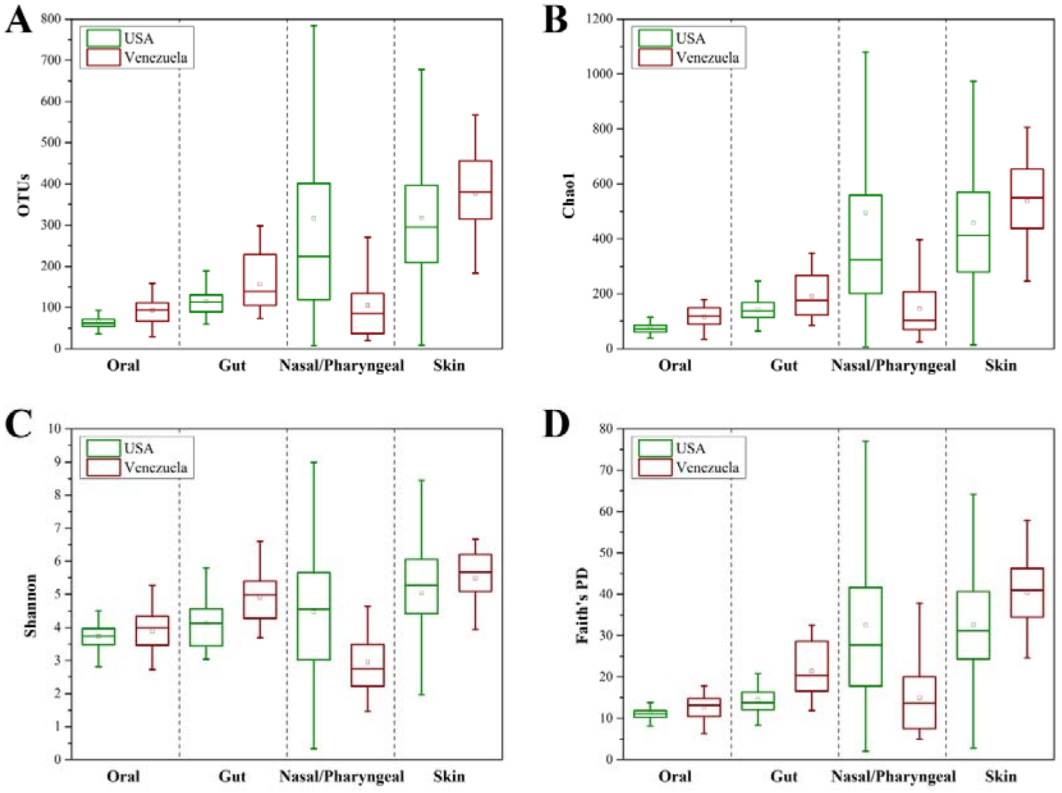


**Supplementary Fig. 3. Alpha diversity comparisons of microbiomes at four different body parts of oral cavity, gut, nose/pharynx and skin between the two different countries of the United States and Venezuela.** (A) Observed OTUs. (B) Shannon index. (C) Chao1 index. (D) Faith's PD value. There were 96 gut samples, 220 nasal/pharyngeal samples, 270 oral samples, and 235 skin samples in the United States. There were 23 gut samples, 33 nasal/pharyngeal samples, 42 oral samples, and 30 skins samples in Venezuelan.
